# Recall of pre-existing cross-reactive B cell memory following Omicron breakthrough infection

**DOI:** 10.1101/2022.04.01.486726

**Authors:** Chengzi I. Kaku, Alan J. Bergeron, Clas Ahlm, Johan Normark, Mrunal Sakharkar, Mattias N. E. Forsell, Laura M. Walker

## Abstract

Understanding immune responses following SARS-CoV-2 breakthrough infection will facilitate the development of next-generation vaccines. Here, we profiled spike (S)-specific B cell responses following Omicron/BA.1 infection in mRNA-vaccinated donors. The acute antibody response was characterized by high levels of somatic hypermutation (SHM) and a bias toward recognition of ancestral SARS-CoV-2 strains, suggesting the early activation of vaccine-induced memory B cells (MBCs). BA.1 breakthrough infection induced a shift in B cell immunodominance hierarchy from the S2 subunit toward the receptor binding domain (RBD). A large proportion of RBD-directed neutralizing antibodies isolated from BA.1 breakthrough infection donors displayed convergent sequence features and broadly recognized SARS-CoV-2 variants of concern (VOCs). Together, these findings provide fundamental insights into the role of pre-existing immunity in shaping the B cell response to heterologous SARS-CoV-2 variant exposure.

**One sentence summary:** BA.1 breakthrough infection activates pre-existing memory B cells with broad activity against SARS-CoV-2 variants.

## Main text

mRNA-based COVID-19 vaccines demonstrated a remarkably high degree of protective efficacy against the original SARS-CoV-2 Wuhan-1 strain in clinical studies (*1, 2*). However, waning vaccine-induced immunity combined with the continued emergence of resistant SARS-CoV-2 variants has significantly undermined vaccine effectiveness (*3–5*). In particular, the recently emerged Omicron variant (B.1.1.529/BA.1) and its sub-lineages (e.g. BA1.1 and BA.2) display a striking degree of antibody evasion, thus severely limiting vaccine efficacy against this VOC and allowing it to rapidly displace Delta and drive a global surge in COVID-19 caseloads (*6–11*).

Understanding the role of antigenic imprinting in shaping the B cell response to antigenically drifted SARS-CoV-2 variants will be critical for the development of next-generation COVID-19 vaccines. Previous studies have shown that breakthrough infection with Delta or Omicron boosts serum neutralizing activity against both the Wuhan-1 vaccine strain and the infecting variant, suggesting recall of cross-reactive vaccine-induced MBCs (*12–14*). However, the specificities, functions, and genetic features of the antibodies mediating this response remain poorly defined. To address these questions, we investigated S-specific serological and peripheral B cell responses in a cohort of mRNA-vaccinated individuals who had recently experienced BA.1 breakthrough infections.

We recruited seven mRNA (mRNA-1273 or BNT162b2)-vaccinated individuals residing in the Northeastern region of the United States who experienced SARS-CoV-2 breakthrough infections between December 30, 2021 and Jan 19, 2022 (Table S1). All donors tested positive for SARS-CoV-2 by RT-PCR and experienced asymptomatic or mild disease. While we were unable to obtain viral samples for genome sequencing, SARS-CoV-2 variant surveillance data indicates that the BA.1 variant accounted for the vast majority of infections in the United States Northeast during this time period (Fig. S1). Breakthrough infections occurred 5-11 months after the second dose of an mRNA vaccine in four donors and one month after a third mRNA dose in three donors. To study the acute B cell response following breakthrough infection, we collected serum and peripheral blood mononuclear cell (PBMC) samples 14 to 27 days following PCR-confirmed infection (Fig. 1A).

**Figure 1.**
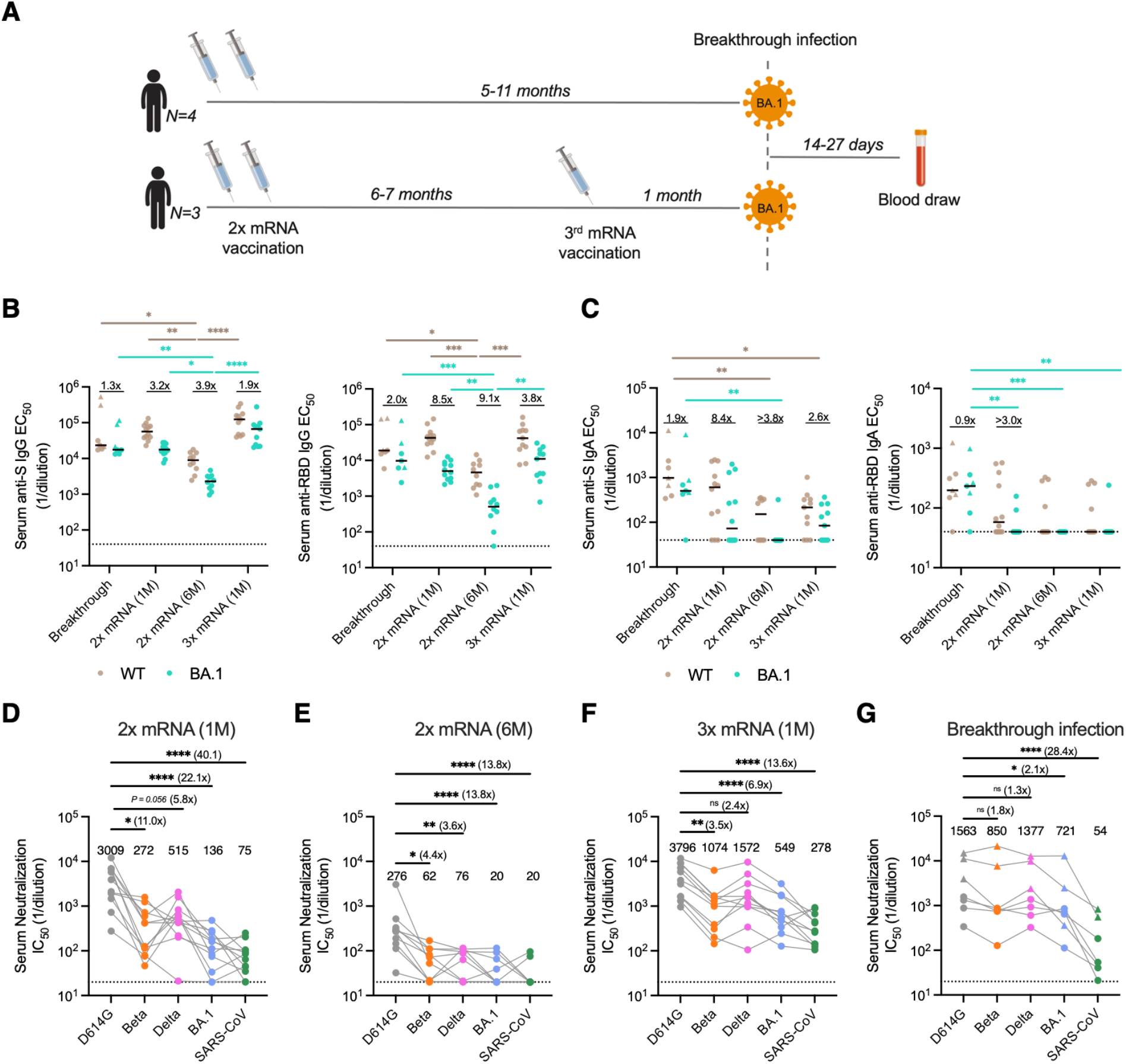
Serum binding and neutralizing activity following BA.1 breakthrough infection. **(A)** Vaccination, infection, and blood draw timelines. **(B-C)** Serum IgG (B) and (C) IgA reactivity with recombinant WT and BA.1 Hexapro-stabilized S proteins (left) and RBDs (right) following BA.1 breakthrough infection. Serum samples from uninfected/vaccinated donors at one month (1M) or six months (6M) following primary vaccination (2x mRNA) or one month following booster mRNA vaccination (3x mRNA) are shown for comparison. The fold change in median EC_50_ against BA.1 relative to D614G is shown above each paired set of measurements. Black bars represent median binding EC_50_ titers. Dotted lines represent the lower limit of detection. **(D-G)** Serum neutralizing activity against SARS-CoV-2 D614G, Beta, Delta, and BA.1 and SARS-CoV (D) one month after primary mRNA vaccination (n=12), (E) six months after primary mRNA vaccination (n=10), (F) one month after booster mRNA vaccination (n=11), and (G) 14 to 27 days after BA.1 breakthrough infection (n=7), as measured using a murine leukemia virus (MLV)-based pseudovirus neutralization assay. Plotted values represent serum neutralizing IC_50_ titers and values shown above the data points indicate the median IC_50_ titer. The fold change in IC_50_ titer for each virus relative to D614G is shown in parentheses. Donors infected after primary mRNA vaccination are shown as circles and those infected after booster mRNA vaccination are shown as triangles. Statistical comparisons were determined by (B-C) two-sided Kruskal-Wallis test with Dunn’s multiple comparisons or (D) Friedman’s test with multiple comparisons. 1M, one month; 6M, six months; EC_50_, 50% effective concentration; IC_50_, 50% inhibitory concentration; WT, wild type. *P < 0.05, **P < 0.01, ***P < 0.001, ****P < 0.0001.

We evaluated serum IgG and IgA responses to recombinant prefusion-stabilized Wuhan-1/wild type (WT) and BA.1 S proteins and RBD subunits following breakthrough infection. For comparison, we also assessed serum antibody responses in a separate cohort of previously uninfected individuals who had received a second dose of an mRNA vaccine at either one- or six-months prior to sampling or a third mRNA booster dose one month prior to sampling (Table S2). Donors who experienced BA.1 breakthrough infection exhibited similar (within two-fold) serum IgG binding titers to BA.1 and WT S and RBD (Fig. 1B). In contrast, uninfected/mRNA vaccinated donors displayed two- to four-fold reduced serum IgG binding to BA.1 relative to WT S and four- to nine-fold lower serum IgG binding titers to the BA.1 RBD relative to WT (Fig. 1B). Furthermore, breakthrough infection donors exhibited significantly higher serum IgA antibody titers to both WT and BA.1 RBDs relative to uninfected/vaccinated donors (Fig. 1C). Thus, BA.1 breakthrough infection induces serum IgG and IgA binding responses to both WT and BA.1 S antigens in previously vaccinated individuals.

Next, we assessed the samples for serum neutralizing activity against an ancestral SARS-CoV-2 strain (D614G), as well as BA.1, Delta, and Beta VOCs using an MLV-based pseudovirus assay. Consistent with prior studies, serum samples obtained from uninfected/vaccinated donors showed 3.5- to 11-fold and 7- to 22-fold lower neutralizing titers against Beta and BA.1, respectively, relative to D614G (Fig. 1D-F). In contrast, serum samples from BA.1 breakthrough infection donors displayed similar (within two-fold) neutralizing titers against D614G and all VOCs tested, suggesting that BA.1 breakthrough infection broadens serum neutralizing antibody responses (Fig. 1G). To determine whether this breadth of activity extended to more divergent sarbecoviruses, we also tested the serum samples for neutralizing activity against SARS-CoV. We observed similar SARS-CoV neutralizing titers in BA.1 breakthrough donors and uninfected/vaccinated individuals, suggesting that the breadth of serum reactivity induced by BA.1 breakthrough infection is likely limited to variants of SARS-CoV-2 and not more antigenically diverse sarbecoviruses (Fig. 1D-G).

We next evaluated the magnitude and cross-reactivity of the peripheral RBD-specific B cell response following BA.1 breakthrough infection. Although we observed higher serum neutralizing titers to BA.1 in breakthrough donors relative to uninfected/mRNA vaccinated individuals, the two cohorts showed similar frequencies of WT- and BA.1-RBD-reactive IgG^+^ B cells (Fig. 2A and Fig. S2A). The limited magnitude of the circulating IgG^+^ B cell response following BA.1 breakthrough infection may be due to the restriction of antigen to the upper respiratory tract during mild and asymptomatic infections. To address this, we also compared the frequencies of RBD-specific IgA^+^ B cells in breakthrough donors and uninfected/vaccinated individuals. In uninfected/vaccinated donors, WT and BA.1 RBD-reactive B cells represented 0.04-0.087% and 0-0.015% of total IgA^+^ B cells, respectively (Fig. 2B). In contrast, breakthrough infection donors mounted significantly higher magnitude IgA responses to the RBD, with BA.1 RBD-specific IgA^+^ B cells accounting for 0.025-0.4% (median=0.069%) of the total IgA^+^ B cell population (Fig. 2B). We conclude that BA.1 breakthrough infection induces similar IgG^+^ B cell responses and higher magnitude IgA^+^ B cell responses to BA.1 RBD antigens relative to mRNA vaccination.

**Figure 2.**
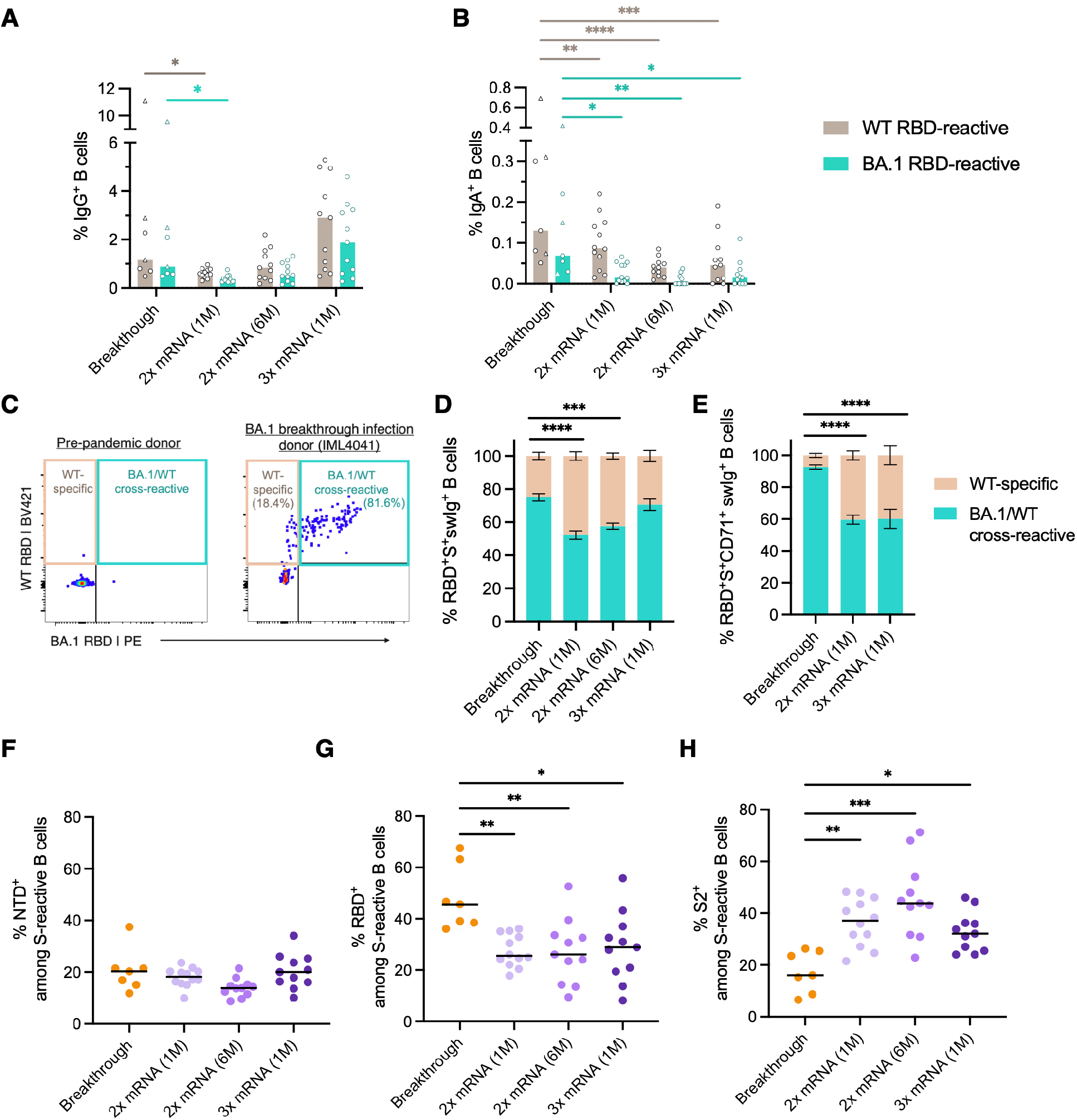
SARS-CoV-2 S-specific B cell responses induced by BA.1 breakthrough infection. **(A-B)** Frequency of circulating B cells that recognize recombinant WT and BA.1 RBDs among (A) IgG^+^ and (B) IgA^+^ B cells in BA.1 breakthrough infection donors and uninfected/vaccinated donors at one- or six-months following primary vaccination (2x mRNA) or one month following booster mRNA vaccination (3x mRNA), as measured by flow cytometry. Bars indicate median frequencies. Donors with breakthrough infections occurring after primary mRNA vaccination are shown as circles and those infected after booster mRNA vaccination are shown as triangles. **(C)** Representative fluorescence-activated cell sorting (FACS) gating strategy used to identify RBD-directed B cells that are WT-specific or WT/BA.1 cross-reactive, shown for a pre-pandemic donor and a breakthrough infection donor. Percentages of WT-specific (tan) and WT/BA.1 cross-reactive (teal) B cells out of all RBD-reactive cells are shown in parentheses. **(D-E)** Mean proportions of RBD-reactive B cells that bind WT and/or BA.1 RBD among (D) total S^+^swIg^+^ B cells or (E) S^+^ swIg^+^ CD71^+^B cells. Error bars represent standard errors of the mean. Samples collected six months following mRNA vaccination were excluded from this analysis due to low numbers of RBD-specific CD71^+^ cells at this time point. **(F-H)** Percentage of S-reactive swIg^+^ B cells that target the (F) NTD, (G) RBD, and (H) Hexapro-stabilized S2 subunits. Black bars represent median percentages. For breakthrough infection donors, this analysis was restricted to S^+^swIg^+^CD71^+^B cells to capture the activated response. Statistical comparisons were determined by (A-B) two-way ANOVA with subsequent Dunnett’s multiple comparisons test or (D-H) two-sided Kruskal-Wallis test with Dunn’s multiple comparisons. 1M, one month; 6M, six months; swIg, class-switched immunoglobulin; WT, wild type. *P < 0.05, **P < 0.01, ***P < 0.001, ****P < 0.0001.

To investigate the impact of pre-existing vaccine-induced immunity on the B cell response to BA.1 breakthrough infection, we enumerated B cells that displayed WT/BA.1 RBD cross-reactivity in BA.1 breakthrough donors and uninfected/mRNA vaccinated individuals (Fig. 2C, Fig. S2A). At one month post primary mRNA vaccination, only 48% of total RBD-directed B cells displayed cross-reactivity with BA.1 (Fig. 2D). The proportion of WT/BA.1 RBD cross-reactive B cells increased to 57% at 6-months post-primary vaccination and to 70% following mRNA booster immunization, consistent with the evolution of anti-SARS-CoV-2 antibody breadth over time (*15, 16*) (Fig. 2D). Following breakthrough infection, BA.1/WT RBD cross-reactive B cells constituted 65-83% of total anti-RBD B cells and the remaining 17-35% bound only to the WT probe (Fig. 2D). Because WT RBD-specific B cells may represent resting MBCs induced by vaccination but not activated by BA.1 infection, we also restricted this analysis to B cells expressing the activation marker CD71 (Fig. S2B). Among recently activated B cells, 87-98% of total RBD-reactive clones displayed BA.1/WT cross-reactivity, whereas the proportion of cross-reactive B cells remained unchanged in uninfected/mRNA vaccinated individuals (Fig. 2E). We were unable to detect BA.1-specific B cells in any donors following BA.1 breakthrough infection, suggesting limited induction of *de novo* B cell responses at this time point. We conclude that BA.1 breakthrough infection preferentially activates B cells that display cross-reactivity with both BA.1 and the original Wuhan-1 vaccine strain.

We next evaluated whether BA.1 breakthrough infection modifies the immunodominance hierarchy of B cells targeting each subdomain with the S trimer. To calculate the proportion of full-length S-reactive B cells targeting each subdomain, we stained B cells with differentially labeled tetramers of full-length S, RBD, NTD, and prefusion-stabilized S2 (Fig. S2C). In the uninfected/vaccinated cohort, class-switched B cells targeting the NTD, RBD, and S2 subdomains comprised 18%, 25%, and 37% of the total S-directed response, respectively, and these proportions remained largely unchanged at six months post-vaccination and after mRNA booster immunization (Fig. 2F-H). In contrast, we observed significantly higher proportions of RBD-directed B cells among donors with breakthrough infection, ranging from 35-63% (median=46%) of the total activated (CD71^+^) B cell response to S (Fig 2G). Furthermore, S2-reactive B cells comprised a smaller fraction (median=16%) of the S-specific response in breakthrough donors relative to uninfected/mRNA vaccinated individuals (Fig 2H). This altered immunodominance pattern was observed in donors experiencing BA.1 breakthrough infection following both second and third dose mRNA vaccination (Fig. S3). In summary, BA.1 breakthrough infection appears to re-direct B cell immunodominance hierarchy from the S2 subunit to the RBD.

To characterize the molecular features of anti-RBD antibodies elicited by BA.1 breakthrough infection, we single-cell sorted 410 class-switched RBD^+^ B cells from five breakthrough infection donors and expressed 317 natively paired antibodies as full-length IgGs (32 to 102 antibodies per donor) (Fig. S4). Despite sorting with a mixture of WT and BA.1 RBDs, the vast majority of IgGs displayed BA.1 RBD reactivity (92-96%), providing strong evidence that these donors experienced BA.1 infections (Fig. 3A). In addition, index sorting analysis revealed that all antibodies derived from CD71^+^B cells recognized BA.1, suggesting that the WT-specific antibodies likely originated from resting MBCs elicited by vaccination (Fig. S5).

**Figure 3.**
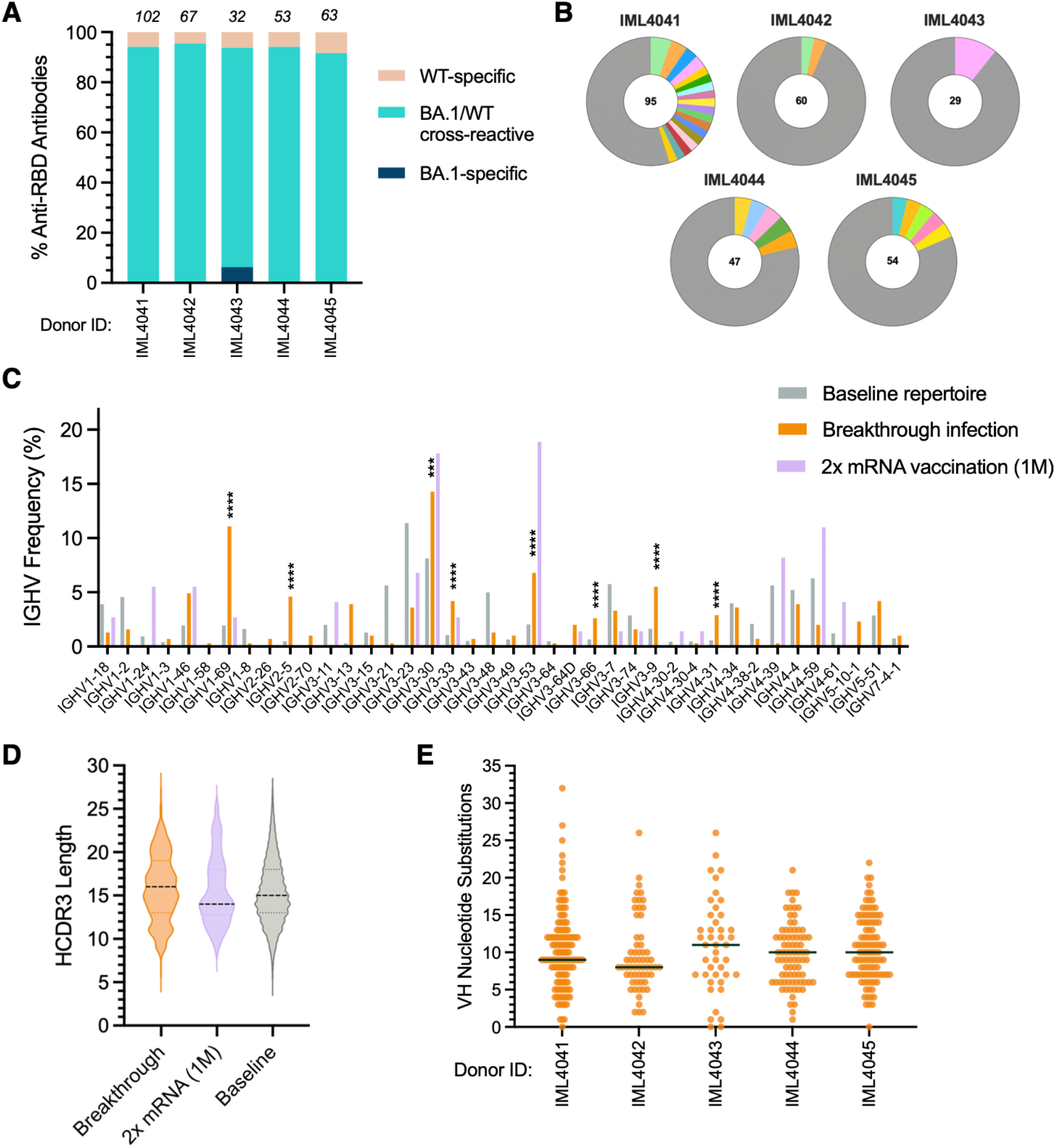
Sequence features of RBD-directed monoclonal antibodies isolated from BA.1 breakthrough infection donors. **(A)** Proportion of antibodies that bind recombinant WT and/or BA.1 RBD antigens from each donor, as determined by biolayer interferometry (BLI). The number of antibodies isolated from each donor is indicated at the top of each bar. **(B)** Clonal lineage analysis. Clonally expanded lineages are represented as colored slices proportional to the lineage size, and unique clones are combined and shown as a single grey segment. The total number of antibodies is shown in the center of each pie. **(C)** Germline IGHV gene usage frequencies among anti-RBD antibodies derived from breakthrough infection donors. Germline gene frequency distributions among anti-RBD antibodies isolated from mRNA vaccinated donors and unselected (baseline) memory B cell repertoires are included for reference (*17*). **(D)** Distribution of HCDR3 amino acid lengths in BA.1-reactive antibodies. The dotted black line represents the median, and the lower and upper lines represent the first and third quartile, respectively. **(E)** Number of VH nucleotide substitutions in antibodies isolated from each breakthrough infection donor, with medians shown by black bars. Statistical significance was determined by Fisher’s exact test compared to the baseline repertoire. 1M, one month; CDR, complementarity determining region; IGHV, immunoglobulin heavy variable domain; VH, variable heavy chain; WT, wild type. ***P < 0.001, ****P < 0.0001.

Sequence analysis revealed that the BA.1 RBD-reactive antibodies displayed a relatively high level of clonal diversity, with 7-45% belonging to expanded clonal lineages (Fig. 3B). We observed a significant over-representation of heavy chain germline genes IGHV3-53, 3-66, 3-30, 1-69, 3-9, and 4-31 in BA.1 breakthrough infection repertoires relative to the baseline human repertoire (*17*) (Fig. 3C). While IGHV3-53, 3-66, and 3-30 germline gene families have also been shown to be over-represented in the antibody response to ancestral SARS-CoV-2 strains, IGHV1-69, 3-9, and 4-31 appear to be unique to the BA.1 breakthrough response (*18*) (Fig. 3C). The BA.1 RBD-reactive antibodies displayed a similar HCDR3 length distribution compared to the baseline repertoire (Fig. 3D). Ninety-five to 100% of antibodies derived from each donor contained somatic mutations, with median SHM levels ranging from 8 to 11 nucleotide substitutions in VH, supporting a memory B cell origin (Fig. 3E). We conclude that the early B cell response to breakthrough infection is dominated by highly mutated clones that cross-react with both WT and BA.1 RBDs.

To further evaluate the binding properties of the BA.1 RBD-reactive antibodies, we measured their monovalent binding affinities for SARS-CoV-2 D614G, BA.1, BA.2, Beta, and Delta RBDs and the SARS-CoV RBD. The majority (204/293) of RBD-directed antibodies bound with high affinity (K_D_<10 nM) to both BA.1 and WT RBDs, supporting selection from an affinity matured B cell population (Fig. 4A). However, approximately 70% of the antibodies displayed higher affinity binding (>2-fold) to the WT RBD relative to BA.1, providing strong evidence for the re-activation of pre-existing vaccine-induced MBCs following BA.1 breakthrough infection (Fig. 4A). In contrast to vaccine-induced anti-RBD antibodies, which often show reduced activity against the Beta VOC, only a minority (<5%) of antibodies derived from BA.1 breakthrough donors displayed loss of binding to Beta relative to WT (*10, 16*) (Fig. 4B). This difference in antibody binding cross-reactivity is likely due to the presence of shared mutations within the Beta and BA.1 RBDs (E484K/A, K417N, and N501Y). Overall, 82% (241/293) of anti-RBD antibodies isolated from breakthrough infection donors displayed monovalent binding to WT, Beta, Delta, BA.1, and BA.2 RBDs, suggesting that BA.1 breakthrough infection activates a large proportion of B cells with broad SARS-CoV-2 VOC recognition (Fig. 4B). Consistent with the weak serum neutralizing activity observed against SARS-CoV, less than 10% of RBD-targeting antibodies exhibited detectable monovalent binding to the SARS-CoV RBD (Fig. 4B). Thus, BA.1 breakthrough infection appears to preferentially expand B cells targeting epitopes that are conserved across SARS-CoV-2 variants but not more antigenically divergent sarbecoviruses.

**Figure 4.**
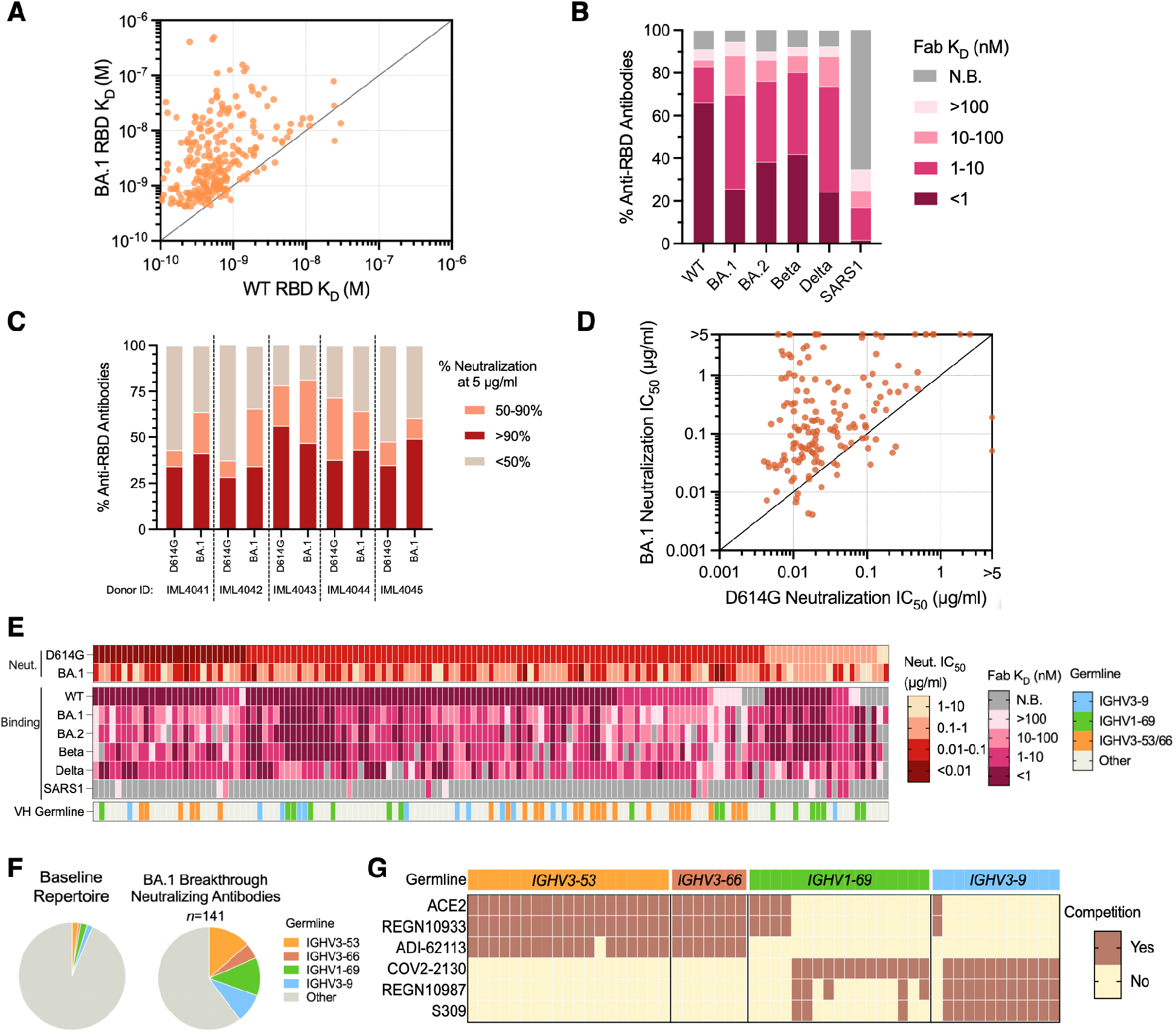
Binding and neutralization properties of anti-RBD antibodies isolated following BA.1 breakthrough infection. **(A)** Fab binding affinities for recombinant WT and BA.1 RBD antigens, as measured by BLI. Fabs with no detectable binding activity or with binding kinetics that could not be fit to a 1:1 binding model were excluded. **(B)** Proportion of BA.1-reactive antibodies with the indicated binding affinities for SARS-CoV-2 VOC RBDs and SARS-CoV RBD, as measured by BLI. Antibodies with weak binding affinities that could not be fit to a 1:1 binding model are shown as >100 nM. **(C)** Proportion of antibodies from each donor with the indicated levels of neutralizing activity against MLV-SARS-CoV-2 D614G and BA.1 at a concentration of 5 μg/ml. **(D)** MLV-SARS-CoV-2 D614G and BA.1 neutralization IC_50_s for antibodies that displayed >90% neutralization against D614G and/or BA.1 at a concentration of 5 μg/ml. **(E)** Heatmap showing neutralization potency and binding breadth of BA.1 neutralizing antibodies. The bottom bar shows convergent IGHV germline gene families. **(F)** Pie charts showing convergent germline gene usage among BA.1-neutralizing antibodies (right) compared to the baseline human antibody repertoires (left) (*17*). **(G)** Competitive binding profiles of BA.1 neutralizing antibodies utilizing convergent IGHV germline genes, as determined by BLI sandwich competition assay using ACE2 and the indicated comparator antibodies. Fab, antigen binding fragment; IC_50_, 50% inhibitory concentration; IGHV, immunoglobulin gene heavy variable; K_D_, equilibrium dissociation constant; N.B., non-binding; WT, wild type.

Next, we screened the BA.1 RBD-reactive antibodies for neutralizing activity against D614G and BA.1. Twenty-eight to 56% and 34-49% of antibodies from each donor displayed >90% neutralizing activity against D614G and BA.1, respectively, at a concentration of 5 μg/ml (Fig. 4C). Titration of the neutralizing antibodies against D614G and BA.1 revealed that 45% (64/141) potently neutralize both viruses with IC_50_s less than 0.1 μg/ml (Fig. 4D). Consistent with their overall increased binding affinities for the WT RBD relative to BA.1, most of the neutralizing antibodies (78%) displayed higher potency against D614G compared to BA.1 (Fig. 4D). Notably, a large proportion of BA.1 neutralizing antibodies also displayed cross-reactivity with Delta (79%), Beta (90%), and BA.2 (86%) RBDs with affinities within 10-fold of BA.1 (Fig. 4E). A limited number of these VOC cross-reactive antibodies (5 out of 141) also neutralized SARS-CoV, with IC_50_s ranging from 0.039-0.35 μg/ml (Fig. S6). We conclude that BA.1 breakthrough infection elicits RBD-directed antibodies with broad activity against SARS-CoV-2 VOCs.

Previous studies have defined several “public” classes of neutralizing antibodies (Class 1-4) induced by SARS-CoV-2 infection and vaccination (*19, 20*). To determine whether BA.1 breakthrough infection also elicits recurrent neutralizing antibody responses, we analyzed the sequence and binding features of the BA.1 neutralizing antibodies. Over 40% of all BA.1 neutralizing antibodies utilized one of three VH germline genes (IGHV3-53/66, IGHV1-69, and IGHV3-9) (Fig. 4F and Fig. S7). Similar to previously described IGHV3-53/66 antibodies isolated from mRNA-vaccinated individuals, the BA.1 neutralizing IGHV3-53/66 antibodies possessed short HCDR3s (11-12 residues) and displayed competitive binding with ACE2, the class 1 mAb REGN10933, and the COVA1-16-like class 4 mAb ADI-62113 (*21*) (Fig. 4G and Fig. S8). However, unlike vaccine-induced IGHV3-53/66 antibodies, which generally lack activity against SARS-CoV-2 variants containing substitutions at position K417 (e.g. Beta, Gamma, and BA.1), breakthrough infection-derived IGHV3-53/66 antibodies displayed broad reactivity with all VOCs tested and potently neutralized both D614G and BA.1 pseudoviruses (median IC_50_ = 0.016 and 0.051 μg/ml, respectively) (*22, 23*) (Fig. 4E and Fig. S9). Thus, these IGHV3-53/66-utilizing antibodies appear to recognize an antigenic site that is overlapping but distinct from previously described IGHV3-53/66 antibodies induced by infection and vaccination.

Neutralizing antibodies utilizing the IGHV1-69 and IGHV3-9 germline genes also broadly recognized SARS-CoV-2 variants, including BA.2 (Fig. 4E). In contrast to IGHV3-53/66, these germline genes have not been shown to be over-represented among RBD-directed antibodies identified from mRNA-vaccinated donors (Fig. 3C). Antibodies utilizing the IGHV1-69 germline gene segregated into two groups, one comprised of antibodies that targeted an ACE2- and REGN10933-competitive region and the other containing antibodies that recognized a non-ACE2 competitive site overlapping the COV2-2130 (class 3) epitope (Fig. 4G). Notably, >80% of non-ACE2 competitive clones utilized the light chain IGLV1-40 gene and displayed highly similar LCDR3 sequences, suggesting a convergent mode of recognition (Fig. S10). Finally, 12/13 IGHV3-9 antibodies recognized an epitope outside of the ACE2 binding site and competed with all three class 3 antibodies tested (S309, REGN10987, and COV2-2130), suggesting a binding mode distinct from the IGHV1-69 antibodies (Fig. 4G). Taken together, BA.1 breakthough infection elicits multiple recurrent classes of anti-RBD antibodies with broad SARS-CoV-2 VOC reactivity.

A deep understanding of how pre-existing SARS-CoV-2 immunity shapes the B cell response to heterologous variant exposure will be important for the development of variant-based booster vaccines. Here we demonstrate that the acute B cell response to BA.1 breakthrough infection is primarily mediated by re-activated vaccine-induced memory B cell clones with broader SARS-CoV-2 VOC cross-reactivity than those elicited by infection or vaccination with ancestral SARS-CoV-2 strains. Although the durability and kinetics of the breakthrough-activated B cell response remain unknown, the induction of cross-reactive responses following BA.1 breakthrough infection suggests that booster immunization with heterologous S proteins may be a promising strategy for the elicitation of broadly neutralizing responses against future emerging VOCs.

Despite the immunodominance of the S2 subunit in the context of primary SARS-CoV-2 infection and vaccination, BA.1 breakthrough infection preferentially boosted cross-reactive antibodies targeting the antigenically variable and immuno-subdominant RBD. The molecular explanation(s) for this shift in B cell immunodominance hierarchy remain to be determined but may be driven by increased serum antibody masking of the conserved S2 subunit relative to the more divergent RBD, resulting in limited S2 epitope accessibility for B cell targeting. Notably, serum antibody feedback via epitope masking has been previously shown to limit B cell responses to immunodominant viral epitopes and permit expansion of subdominant responses (*24*).

Finally, we identified several monoclonal antibodies from BA.1 breakthrough infection donors that display broad activity against all SARS-CoV-2 VOCs described to date as well as SARS-CoV. These antibodies represent promising candidates for therapeutic development and provide a framework for the development of vaccines that induce broadly neutralizing antibody responses.

## Acknowledgements

We thank study nurse Ida-Lisa Persson and personnel at the Clinical Research Center at Umeå University Hospital for support, enrollment of study subjects, and sampling. We acknowledge Linnea Vikström at Umeå University for processing of samples and summary of clinical data. We also thank E. Krauland, J. Nett, and M. Vasquez for helpful comments on the manuscript. All IgGs were sequenced by Adimab’s Molecular Core and produced by the High Throughput Expression group.

## Author contributions

C.I.K. and L.M.W. conceived and designed the study. A.J.B. and M.N.E.F. supervised and performed clinical sample collection and processing. C.A. and J.N. are principal investigators for the clinical trial and supervised sample collection. C.I.K. designed and performed serum and B cell analyses, single B cell sorting, and biolayer interferometry assays. C.I.K. and M.S. performed pseudovirus neutralization assays. C.I.K. and L.M.W. analyzed the data. C.I.K. and L.M.W. wrote the manuscript, and all authors reviewed and edited the paper.

## Data and material availability

Antibody sequences have been deposited in GenBank (accession codes). All other data are available in the manuscript or supplementary materials. IgGs are available from L.M.W. under a material transfer agreement (MTA) from Adagio Therapeutics, Inc.

## Supplementary Materials

Materials and Methods

Figures S1 – S10

Table S1 – S2

References 25 – 26

