## Supplementary material for "Recall of pre-existing cross-reactive B cell memory following Omicron breakthrough infection"

**Materials and Methods**

**Human subjects and blood sample collection.**

Breakthrough infection donors and uninfected two-dose vaccinated donors participated with informed consent under the healthy donor protocol D10083, Immune Monitoring Core (DartLab) Laboratory at Dartmouth-Hitchcock Hospital. Uninfected three-dose vaccinated participants are enrolled in the clinical trial, CoVacc - Immune response to vaccination against Covid-19, an open multicenter phase IV study, was approved by the Swedish Ethics Review Authority (Dnr 2021-00055) and the Medical Products Agency Sweden. The study was registered at European Clinical Trials Database (EUDRACT Number 2021-000683-30) before the first patient was enrolled. Umeå University, Sweden served as trial sponsor and the Clinical Research Center, University Hospital of Northern Sweden was monitoring the study for regulatory compliance. Individuals were included after informed consent and data were stored in accordance with the EU General Data Protection Regulation.

Seven participants with BA.1 breakthrough infection were recruited to participate in this study. SARS-CoV-2 infection was determined by positive results via both RT-PCR from a saliva sample and rapid antigen test from a nasal swab sample. All participants were previously immunized with two- or three-doses of an mRNA vaccine (BNT162b2 or mRNA-1273) and had no documented history of SARS-CoV-2 infection prior to vaccination. Clinical and demographic characteristics of breakthrough infection donors are shown in Table S1. Participants presented to the Dartmouth-Hitchcock Hospital (D-HH) 14 to 27 days after their first SARS-CoV-2 positive test for blood draw. Venous blood was collected using BD Vacutainer® tubes with acid citrate dextrose (ACD), and plasma and PBMCs were isolated using a Ficoll 1077 (Sigma) gradient, washed, and counted with an anti-human CD45 stain on a volumetric flow cytometer. PBMC were frozen in 12.5% human serum and 10% DMSO diluted in RPMI-1040 and stored in liquid nitrogen until use. Plasma was isolated and frozen at -80 ˚C.

A separate cohort of uninfected/mRNA-vaccinated volunteers were recruited for blood sample collection at D-HH (for two-dose mRNA-vaccinated donors) and Umeå University (for three-dose mRNA-vaccinated donors). Samples were collected at one month (n=12) and six months (n=11) following the second mRNA dose or one month following the third mRNA dose (n=11). Demographics for these participants are shown in Table S2. Sample collection and processing methods for two-dose vaccinated individuals are described above. For individuals who received a third mRNA dose, venous blood was collected and PBMCs and plasma isolated in BD EDTA Vacutainer® CPT™ tubes. PBMCs were frozen in 90% fetal calf serum supplemented with 10% DMSO and stored in liquid nitrogen until use. Plasma and serum were stored at -80 ˚C.

**Recombinant SARS-CoV-2 S production.**

To produce prefusion-stabilized WT SARS-CoV-2 HexaPro S, DNA encoding residues 1-1208 of the SARS-CoV-2 spike (Genbank NC NC_045512.2) with substitutions F817P, A892P, A899P, A942P, K986P, V987P, “GSAS” mutations from positions 682-685 and a C-terminal T4 fibritin motif, 8X HisTag and TwinStrepTag (SARS-CoV-2 S-2P) was cloned into a pcDNA3.4 vector. The following mutations were additionally cloned into the Omicron/BA.1 HexaPro S plasmid: A67V, Δ69-70, T95I, G142D, Δ143-145, Δ211, L212I, ins214EPE, G339D, S371L, S373P, S375F, K417N, N440K, G446S, S477N, T478K, E484A, Q493K, G496S, Q498R, N501Y, Y505H, T547K, D614G, H655Y, N679K, P681H, N764K, D796Y, N856K, Q954H, N969K, L981F. Plasmids were transiently transfected into FreeStyle HEK 293F cells (Thermo Fisher) using polyethylenimine following the manufacturer's directions. After one week of culture, the supernatants were harvested, and centrifuged to remove cellular debris. S protein preps were purified by Ni affinity chromatography and followed by size exclusion chromatography using the Superose 6 column (GE Healthcare) before concentrating and freezing at -80 ˚C.

**Serum ELISAs.**

96-well half-area plates (Corning) were coated with the following recombinant antigens at a concentration of 5 µg/ml diluted in PBS: SARS-CoV-2 WT Hexapro-stabilized S, BA.1 Hexapro-stabilized S, WT RBD (Sino Biological, Cat #40592-V08B), BA.1 RBD (Acro Biosystems, Cat #SPD-C522e), WT NTD (Acro Biosystems, Cat #S1D-52H6), BA.1 NTD (Acro Biosystems, Cat #SPD-C522d), and Hexapro-stabilized S2 (Acro Biosystems, Cat #S2N-C52H5) antigens. Following overnight incubation at 4 ˚C, wells were washed with wash buffer (1X PBS with 0.05% Tween-20) and blocked with 75 µl 3% bovine serum albumin (BSA) in 1X PBS for 1 h at 37 °C. Coated wells were subsequently incubated with serial dilutions of human sera ranging from 1:40 to 1:1,310,720 in a solution of 0.1% BSA, 0.01% Tween-20 in 1X PBS for 1 h at 37 °C and then washed three times with wash buffer. To detect antigen-specific IgG and IgA, wells were incubated with either a 1:5000 dilution of anti-human IgG horseradish peroxidase (HRP; Jackson Immunoresearch Laboratories, Cat #109-036-098) or a 1:10,000 dilution of anti-human IgA HRP (Jackson Immunoresearch Laboratories, Cat #109-036-011) in 0.1% BSA, 0.01% Tween-20, 1X PBS for 1 h at 37 ˚C. Plates were then washed three times and developed with 25 µl of room temperature-equilibrated 1-Step™ Ultra TMB Substrate Solution (Thermo Fisher Scientific) for 5 min. The developing reaction was terminated by addition of 25 µl 4 N sulfuric acid. Absorbance was measured at 450 nm using a Spectramax microplate reader (Molecular Devices). Titration curves were fitted via non-linear regression to determine the 50% effective concentration (EC_50_) in GraphPad Prism (version 9.3.1).

**SARS-CoV-2 pseudovirus generation.**

Single-cycle infection pseudoviruses were generated as previously described (*25*). Briefly, HEK293T cells seeded overnight in 6-well tissue culture plates (Corning) were co-transfected with the following plasmids: 1) 0.5 µg of pCDNA3.3 encoding SARS-CoV-2 spike genes with 19-residue C-terminal truncations, 2) 2 µg of MLV-based luciferase reporter gene plasmid (Vector Builder), and 3) 2 µg of MLV gag/pol (Vector Builder). The SARS-CoV-2 Omicron/BA.1 contained the following mutations in relative to Wuhan-1: A67V, Δ69-70, T95I, G142D, Δ143-145, Δ211, L212I, ins214EPE, G339D, S371L, S373P, S375F, K417N, N440K, G446S, S477N, T478K, E484A, Q493K, G496S, Q498R, N501Y, Y505H, T547K, D614G, H655Y, N679K, P681H, N764K, D796Y, N856K, Q954H, N969K, L981F. Plasmids were combined with Lipofectamine 2000 (ThermoFisher Scientific) and transfected following the manufacturer's recommendations. Culture supernatants containing SARS-CoV-2 S-pseudotyped MLV particles were collected 48 h post-transfection, aliquoted, and frozen at -80 °C for neutralization assays.

**Pseudovirus neutralization assay.**

HeLa-hACE2 reporter cells (BPS Bioscience Cat #79958) were seeded overnight at 10,000 cells per well in 96-well tissue culture plates (Corning). Human plasma and serum samples were heat-inactivated at 56 ˚C for 30 min. Next, monoclonal antibodies or heat-inactivated sera were serially diluted in MEM/EBSS media supplemented with 10% FBS with 50 µl of MLV viral stock and incubated for 1 h at 37 ˚C with 5% carbon dioxide. Cell culture media was removed, and cells were washed two times with PBS. The virus-antibody mixture was subsequently added to HeLa-hACE2 cells and incubated for 48 h at 37 ˚C with 5% carbon dioxide. Cells were then lysed with Luciferase Cell Culture Lysis 5× reagent (Promega), and luciferase activity was measured using the Luciferase Assay System (Promega) following manufacturer's protocols. Infectivity was measured as relative luminescence units (RLUs) using a luminometer (Perkin Elmer). The percentage neutralization was calculated as 100*(1–[RLU_sample_– RLU_background_]/[ RLU_isotype control mAb_–RLU_background_]), and the 50% neutralization concentration was interpolated from four-parameter non-linear regression fitted curves in GraphPad Prism (version 9.3.1).

**FACS analysis of SARS-CoV-2 S-specific B cell responses.**

Antigen-specific B cells were detected using recombinant biotinylated antigens tetramerized with fluorophore-conjugated streptavidin (SA). For detection of peripheral B cells that recognize WT and/or BA.1 RBD, 4:1 molar ratios of biotinylated antigens to SA were mixed in the following combinations: WT HexaPro S with SA-AlexaFluor 633 (Invitrogen), BA.1 HexaPro S with SA-AlexaFluor 633 (Invitrogen), WT RBD (Acro Biosystems, Cat #SPD-C82E8) with SA-BV421 (BioLegend), and BA.1 RBD (Acro Biosystems, Cat #SPD-C522e) with SA-phycoerythrin (PE; Invitrogen). For determination of subdomain reactivities within the total S-specific B cell population, antigen tetramers were mixed in the following combinations: WT HexaPro S with SA-AlexaFluor 633 (Invitrogen), BA.1 HexaPro S with SA-AlexaFluor 633 (Invitrogen), WT RBD (Acro Biosystems, Cat #SPD-C82E8) with SA-BV421 (BioLegend), BA.1 RBD (Acro Biosystems, Cat #SPD-C522e) with SA-BV421, WT NTD (Acro Biosystems, Cat #S1D-52H6) with SA-PE, BA.1 NTD (Acro Biosystems, Cat # SPD-C522d) with SA-PE, and HexaPro-stabilized WT S2 (Acro Biosystems, Cat #S2N-C52H5) with SA-BV711 (BD BioSciences). Antigen tetramers were incubated for 30 min at 4 ˚C, followed by quenching of unbound SA sites using 5 µl of 2 µM Pierce biotin (ThermoFisher Scientific). PBMCs were stained with pooled tetramerized antigens (25 nM each) and anti-human antibodies anti-CD19 (PE-Cy7; Biolegend), anti-CD3 (PerCP-Cy5.5; Biolegend), anti-CD8 (PerCP-Cy5.5; Biolegend), anti-CD14 (PerCP-Cy5.5; Invitrogen), and anti-CD16 (PerCP-Cy5.5; Biolegend) diluted in a 1:1 [v/v] mixture of Brilliant Stain Buffer (BD BioSciences) and FACS buffer (2% BSA/1 mM EDTA in 1X PBS) for 15 min on ice. Following one wash, cells were resuspended in a mixture of propidium iodide and anti-human antibodies anti-IgG (BV605; BD Biosciences), anti-IgA (FITC; Abcam), anti-CD27 (BV510; BD Biosciences), and anti-CD71 (APC-Cy7; Biolegend) and incubated for 15 min on ice. After washing two times with FACS buffer, samples were analyzed using a BD FACS Aria II (BD Biosciences).

The proportion of class-switched RBD-specific B cells that reacted with WT and/or BA.1 RBD was calculated by dividing the number of BA.1/WT cross-reactive or WT-specific IgG^+^ and IgA^+^ (swIg^+^) B cells by the total number of RBD^+^ S^+^ swIg^+^ B cells. The proportion of S-reactive B cells that recognized each subdomain (NTD, RBD, or S2) was calculated by dividing the number of IgG^+^ and IgA^+^ (swIg^+^) B cells that recognize both S and the subdomain by the total number of S^+^ swIg^+^ cells.

**Single B cell sorting.**

Biotinylated recombinant WT (Acro Biosystems, Cat #SPD-C82E8) and BA.1 (Acro Biosystems, Cat #SPD-C522e) RBDs were separately mixed with AlexaFluor 633-conjugated SA (SA-633; Invitrogen) and PE-SA (Invitrogen) at a 4:1 molar ratio of antigen to SA for 30 min at 4 ˚C. The four antigen-SA pairs were then pooled to create a mixture of PE- and APC-labelled WT and BA.1 RBD tetramers. PBMCs were stained with tetramerized antigens (25 nM each) and a mixture of anti-CD19 (PE-Cy7; Biolegend), anti-CD20 (BV711; Biolegend) anti-CD3 (PerCP-Cy5.5; Biolegend), anti-CD8 (PerCP-Cy5.5; Biolegend), anti-CD14 (PerCP-Cy5.5; Invitrogen), and anti-CD16 (PerCP-Cy5.5; Biolegend) antibodies diluted in FACS buffer (2% BSA/1 mM EDTA in 1X PBS) for 15 min on ice. Stained cells were centrifuged for 10 min at 400 x *g,* washed once with FACS buffer, and centrifuged again to pellet the cells. Next, cells were resuspended in propidium iodide and anti-human antibodies anti-IgM (BV421; BD Biosciences), anti-IgG (BV605; BD Biosciences), anti-IgA (FITC; Abcam), anti-CD27 (BV510; BD Biosciences), and anti-CD71 (APC-Cy7; Biolegend) diluted in Brilliant Stain Buffer (BD BioSciences) and FACS buffer. Following 15 min of incubation on ice, Cells were washed two times, resuspended in FACS buffer, and analyzed using a BD FACS Aria II (BD Biosciences). Class-switched B cells, defined as CD19^+^CD3^−^CD8^−^CD14^−^CD16^−^PI^−^IgM^−^ and IgG^+^ or IgA^+^, that specifically bound to the WT/BA.1 RBD mixture were single-cell index sorted into 96-well polystyrene microplates (Corning) containing 20 µl lysis buffer per well [5 µl of 5X first strand SSIV cDNA buffer (Invitrogen), 1.25 µl dithiothreitol (Invitrogen), 0.625 µl of NP-40 (Thermo Scientific), 0.25 µl RNaseOUT (Invitrogen), and 12.8 µl dH2O]. Plates were then frozen at -80 ˚C before further downstream processing.

**Amplification and cloning of antibody variable genes.**

Antibody variable gene mRNA transcripts (VH, Vκ, Vλ) were amplified by RT-PCR as described previously (*22*). Briefly, cDNA was synthesized using SuperScript IV enzyme (ThermoFisher Scientific), followed by two rounds of nested PCRs. The second cycle of nested PCR added 40 base pairs of 5’ and 3’ homology to restriction enzyme-digested *S. cerevisiae* expression vectors to enable homologous recombination during transformation. PCR-amplified variable gene DNA was chemically transformed into competent yeast cells via the lithium acetate method and yeast were plated on selective amino acid drop-out agar plates (*26*). Transformed yeast colonies were picked for sequencing and characterization.

**Expression and purification of IgG and Fab molecules.**

Antibodies were expressed as human IgG1 via *S. cerevisiae* cultures, as described previously (*22*). Briefly, yeast cells were grown for IgG expression over 6 days, and the IgG-containing supernatant was subsequently harvested by centrifugation. Antibodies were purified by protein A-affinity chromatography, eluted with a solution of 200 mM acetic acid/50 mM NaCl (pH 3.5). The pH was then neutralized using 1/8^th^ volume of 2 M Hepes (pH 8.0)

Fab fragments were generated by incubating IgG with papain for 2 h at 30 °C. The reaction was terminated using iodoacetamide, and the mixture containing digested Fab and Fc was purified by Protein A agarose to remove Fc fragments and undigested IgG. Fabs present in the flow-through were further purified using CaptureSelect™ IgG-CH1 affinity resin (ThermoFisher Scientific) and eluted from the column using 200 mM acetic acid/50 mM NaCl (pH 3.5). Fab solutions were pH-neutralized using 1/8th volume 2 M Hepes (pH 8.0).

**Binding affinity measurements by biolayer interferometry.**

Binding affinities were measured by biolayer interferometry (BLI) using a FortéBio Octet HTX instrument (Sartorius). All steps were performed at 25 ^o^C and at an orbital shaking speed of 1000 rpm. All reagents were formulated in PBSF buffer (PBS with 0.1% w/v BSA).

Recombinant biotinylated antigens were diluted (100 nM) in PBSF and loaded onto streptavidin biosensors (Sartorius) to a sensor response of 0.6-1.0 nm and then allowed to equilibrate in PBSF for a minimum of 30 min. After a 60 s baseline step in PBSF, antigen-loaded sensors were exposed (180 s) to Fab or IgG fragments (100 nM) and then dipped (180 s) into PBSF to measure any dissociation of the antigen from the biosensor surface. Fab binding data with detectable binding responses (>0.1 nm) were aligned, inter-step corrected (to the association step) and fit to a 1:1 binding model using the FortéBio Data Analysis Software, version 11.1.

**Epitope binning by biolayer interferometry.**

Antibody competition with recombinant human ACE2 and comparator antibodies for binding to SARS-CoV-2 RBD was determined by BLI using a ForteBio Octet HTX (Sartorius). All binding steps were performed at 25 ^o^C and at an orbital shaking speed of 1000 rpm. All reagents were formulated in PBSF (1X PBS with 0.1% w/v BSA). For ACE2 competition experiments, test antibodies (100 nM) were captured onto anti-human IgG capture (AHC) biosensors (Molecular Devices) to a sensor response of 1.0 nm-1.4 nm. IgG-loaded sensors were then soaked (20 min) in an irrelevant IgG1 solution (0.5 mg/ml) to block remaining Fc binding sites, followed by a 30 min incubation in PBSF. To assess any potential cross interactions between sensor-loaded IgG and ACE2, the IgG-loaded and blocked sensors were exposed (90 s) to a 300 nM ACE2 (Sino Biological, Cat# 10108-H08H). Sensors were next allowed to baseline (60 s) before exposing (180 s) to recombinant SARS-CoV-2 RBD (100 nM; Acro Biosystems, Cat # SPD-C52H3) and then exposed (180 s) to ACE2 (300 nM). Increased sensor responses following ACE2 exposure represented a non-ACE2-competitive binding profile, whereas antibodies showing unchanged sensor responses were designated as ACE2-competitive. Antibody competition with comparator antibodies (REGN10933, ADI-62113, COV2-2130, REGN10987, and S309) was performed using the same method as described above but with a different assay orientation: comparator antibodies were captured to anti-human IgG capture biosensors (Molecular Devices) and then exposed to antibodies of interest (300 nM) in solution.

**Supplementary Figures**

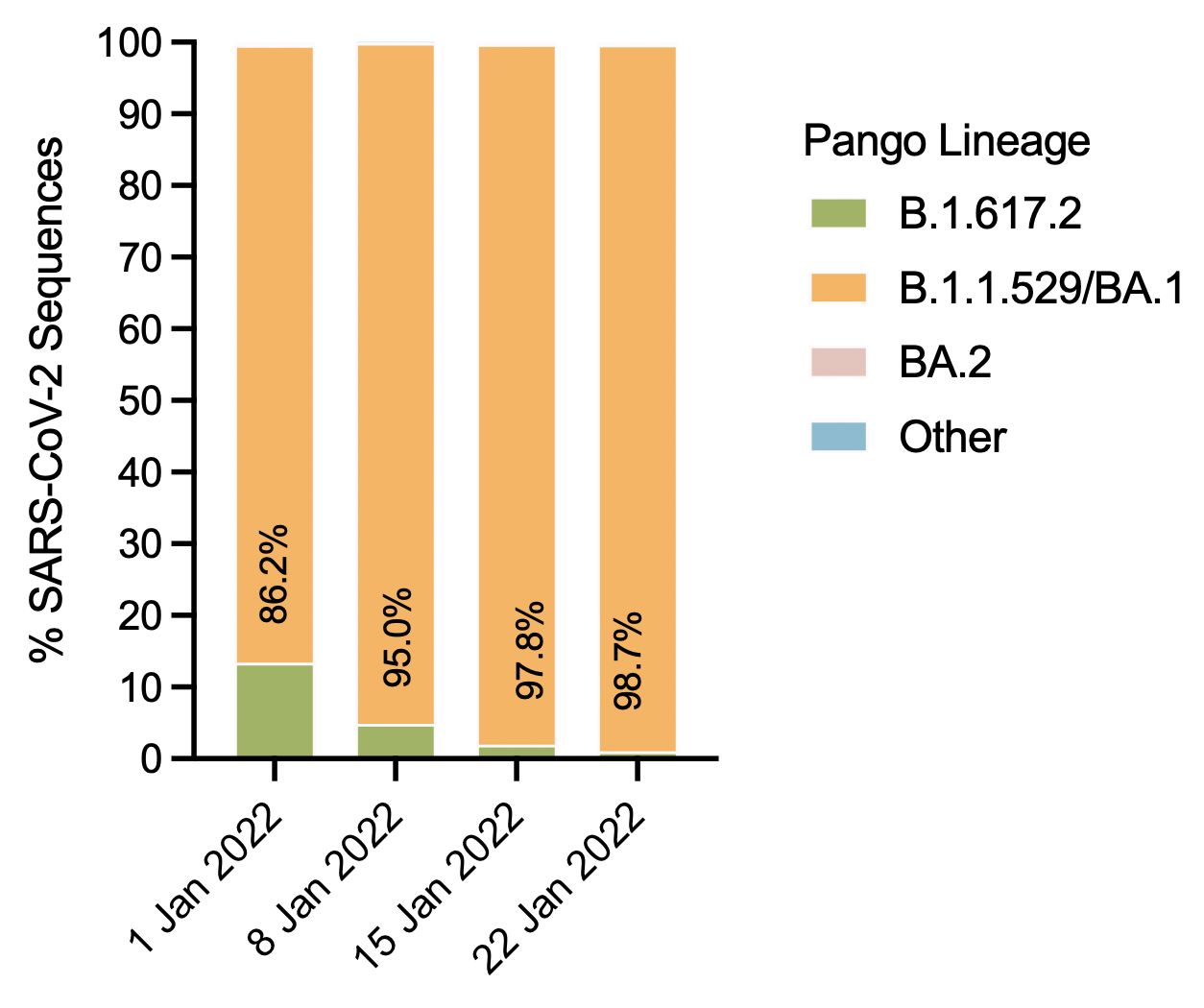

**Fig. S1.** Prevalence of circulating SARS-CoV-2 variants. Genomic sequencing analyses of SARS-CoV-2 infections occurring in the United States CDC Region 1 (Connecticut, Maine, Massachusetts, New Hampshire, Rhode Island, and Vermont) are colored by the indicated Pango lineage. Data was obtained using <https://covid.cdc.gov/covid-data-tracker/#variant-proportions> (accessed March 18, 2022).

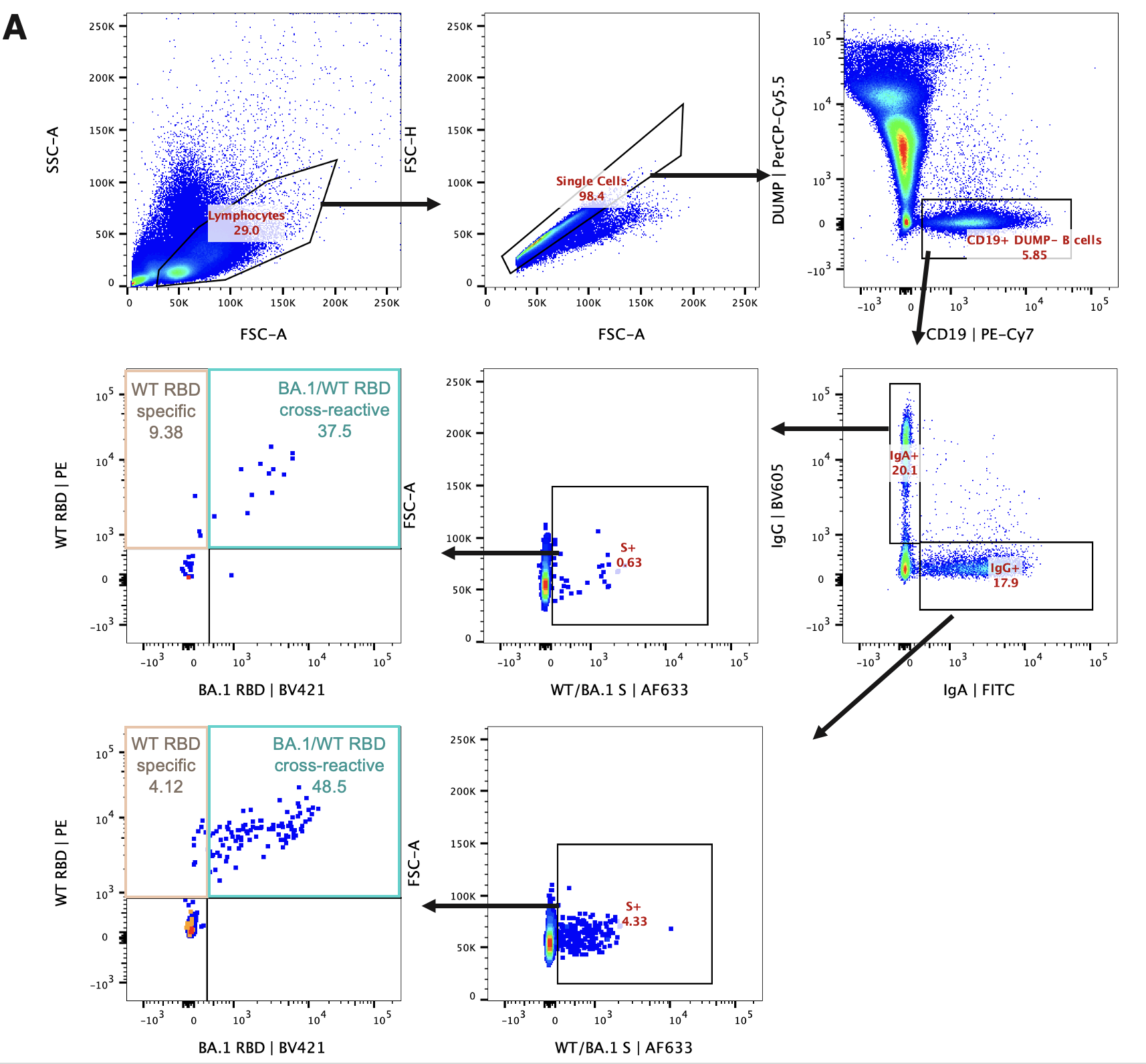

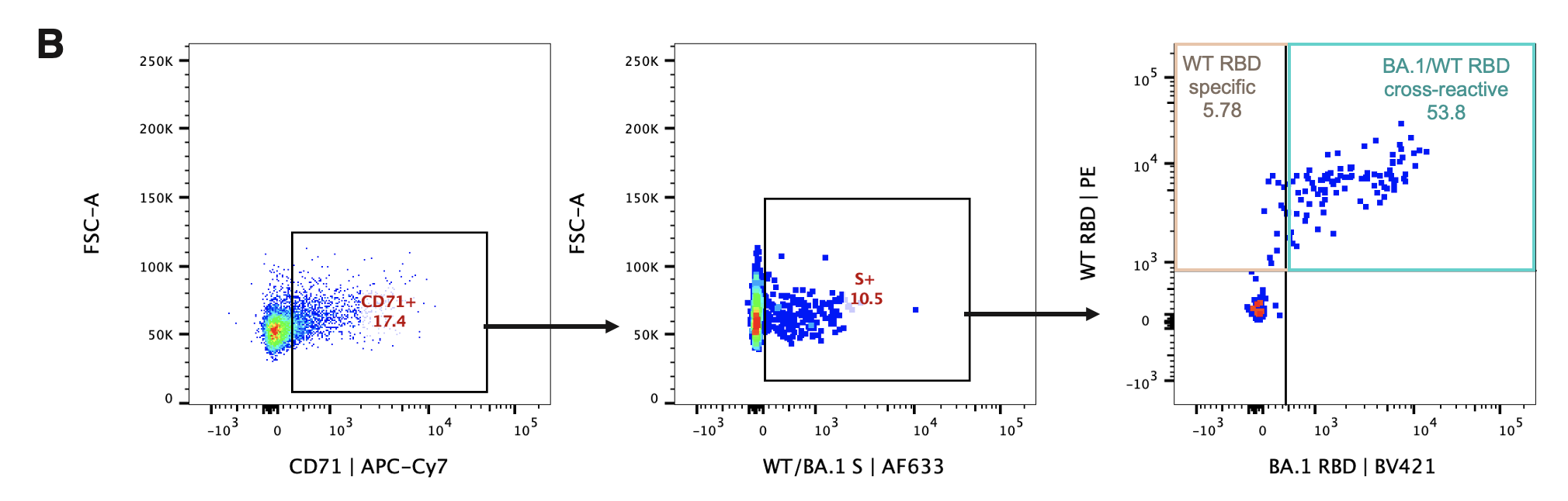

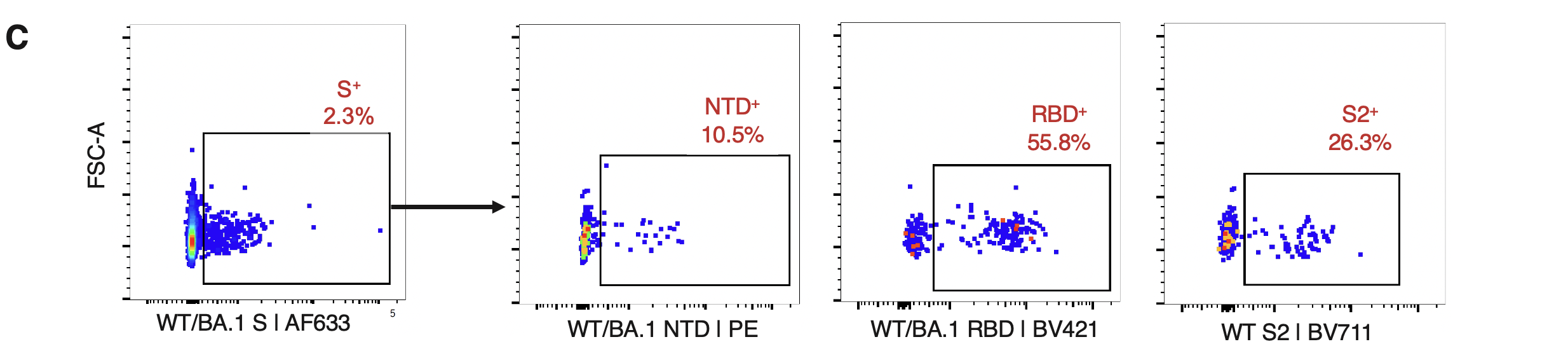

**Fig. S2.** SARS-CoV-2 RBD-specific B cell staining. **(A)** Representative FACS gating strategy to determine frequencies of WT- and BA.1-RBD-reactive B cells among IgG^+^ and IgA^+^ B cells. **(B)** Representative FACS gating strategy to determine the proportion of CD71^+^S^+^RBD^+^ cells that are WT-specific or BA.1/WT cross-reactive. **(C)** Representative FACS gating strategy used to calculate the proportion S-specific B cells directed to the NTD, RBD, and S2 subdomains. FSC-A, forward scatter area; FSC-H, forward scatter height; SSC-A, side scatter area.

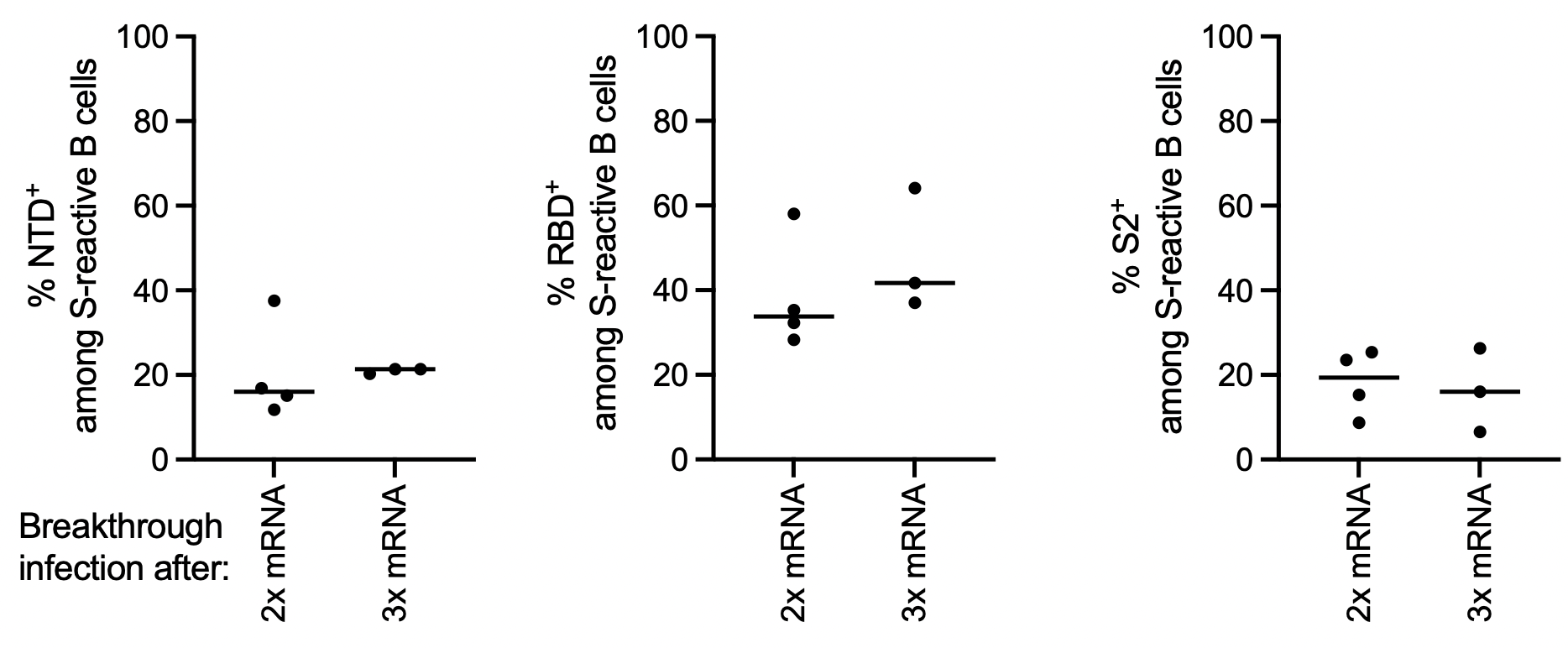

**Fig. S3.** Distribution of subdomain specificities within the S-specific B cell response following BA.1 breakthrough infection. Proportions of S-reactive CD71^+^ swIg^+^ B cells that recognize the NTD (left), RBD (middle), or prefusion-stabilized S2 (right) subdomains in donors who experienced breakthrough infection after either two-dose or three-dose mRNA vaccination, as determined by flow cytometry. Black bars indicate medians.

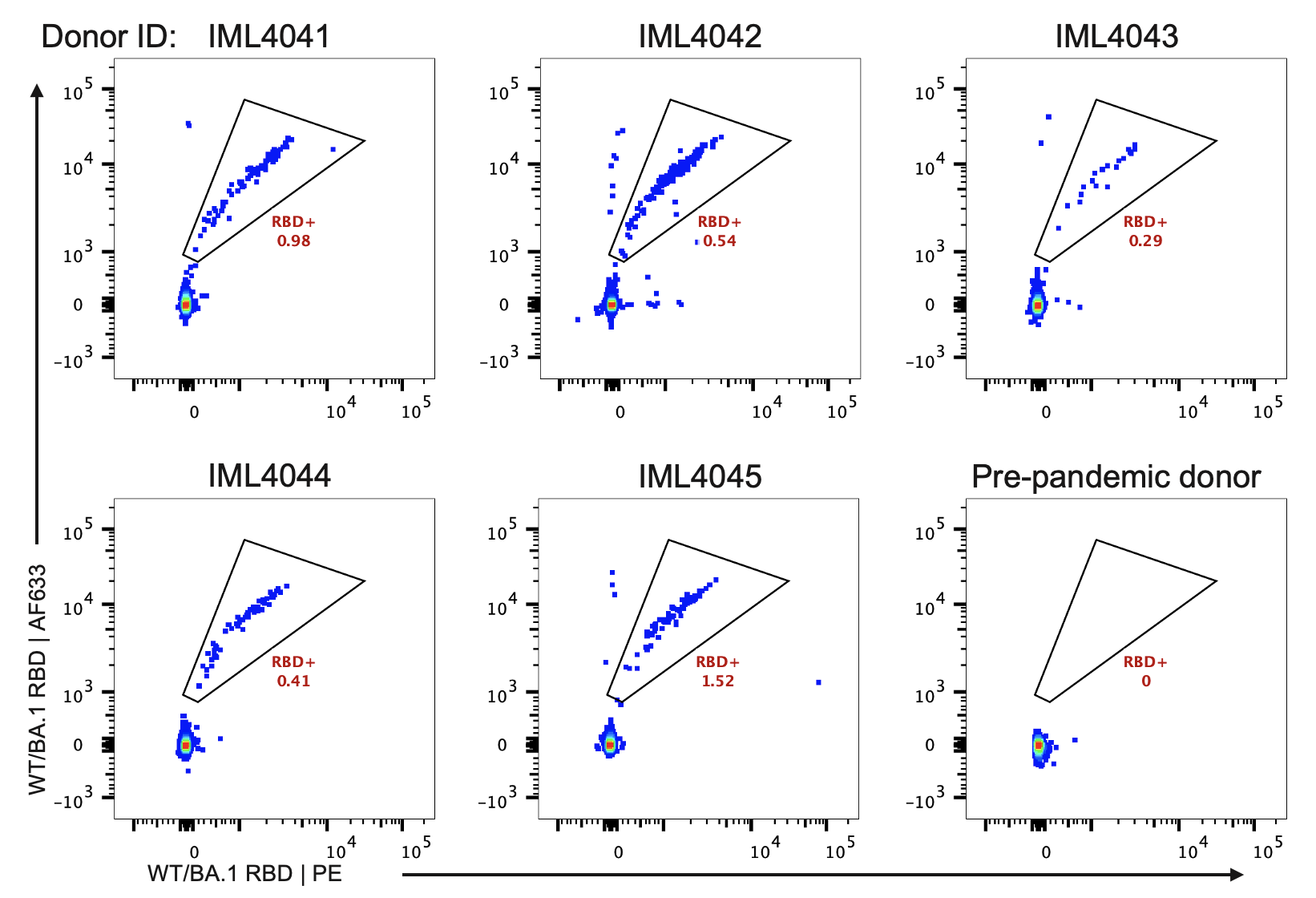

**Fig. S4.** Representative FACS gating strategy for single-cell sorting RBD-specific swIg^+^ B cells. FACS plots shown are gated on swIg^+^ CD19^+^ B cells. A pre-pandemic donor was included as a negative control.

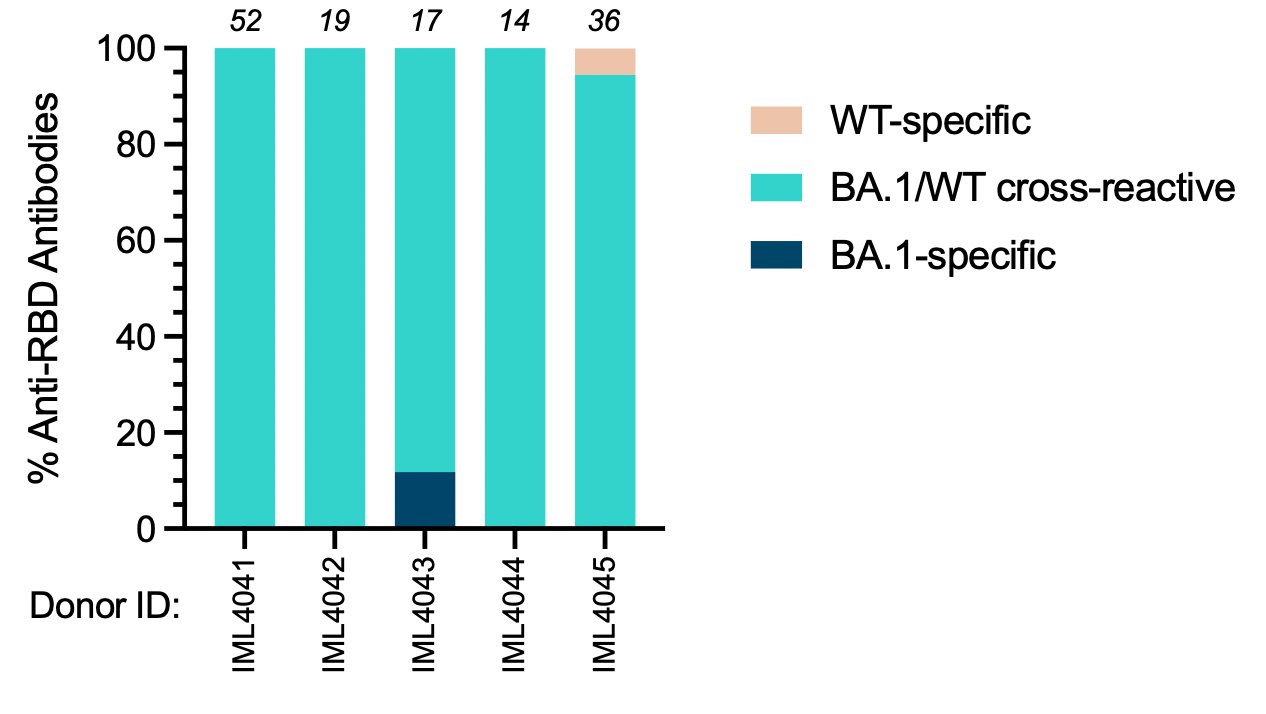

**Fig. S5.** Binding reactivities of breakthrough infection-derived antibodies isolated from CD71^+^ B cells. The number at the top of each bar represents the total number of antibodies.

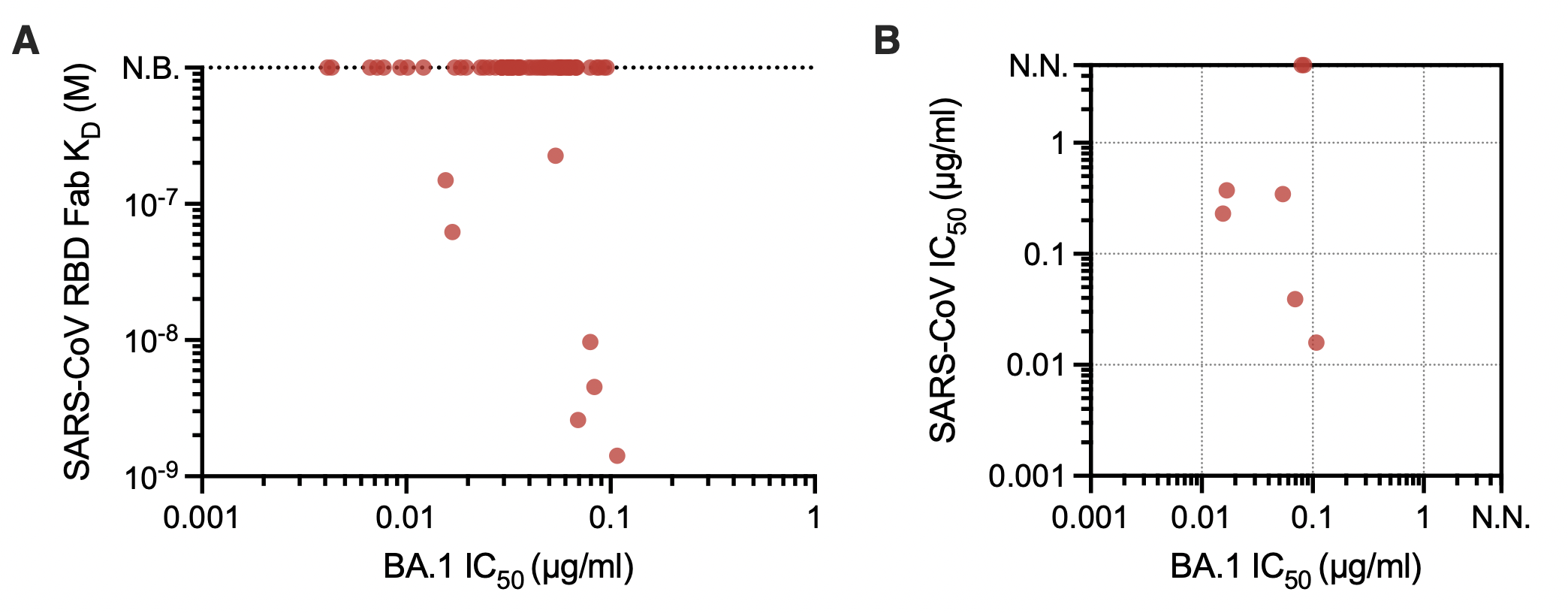

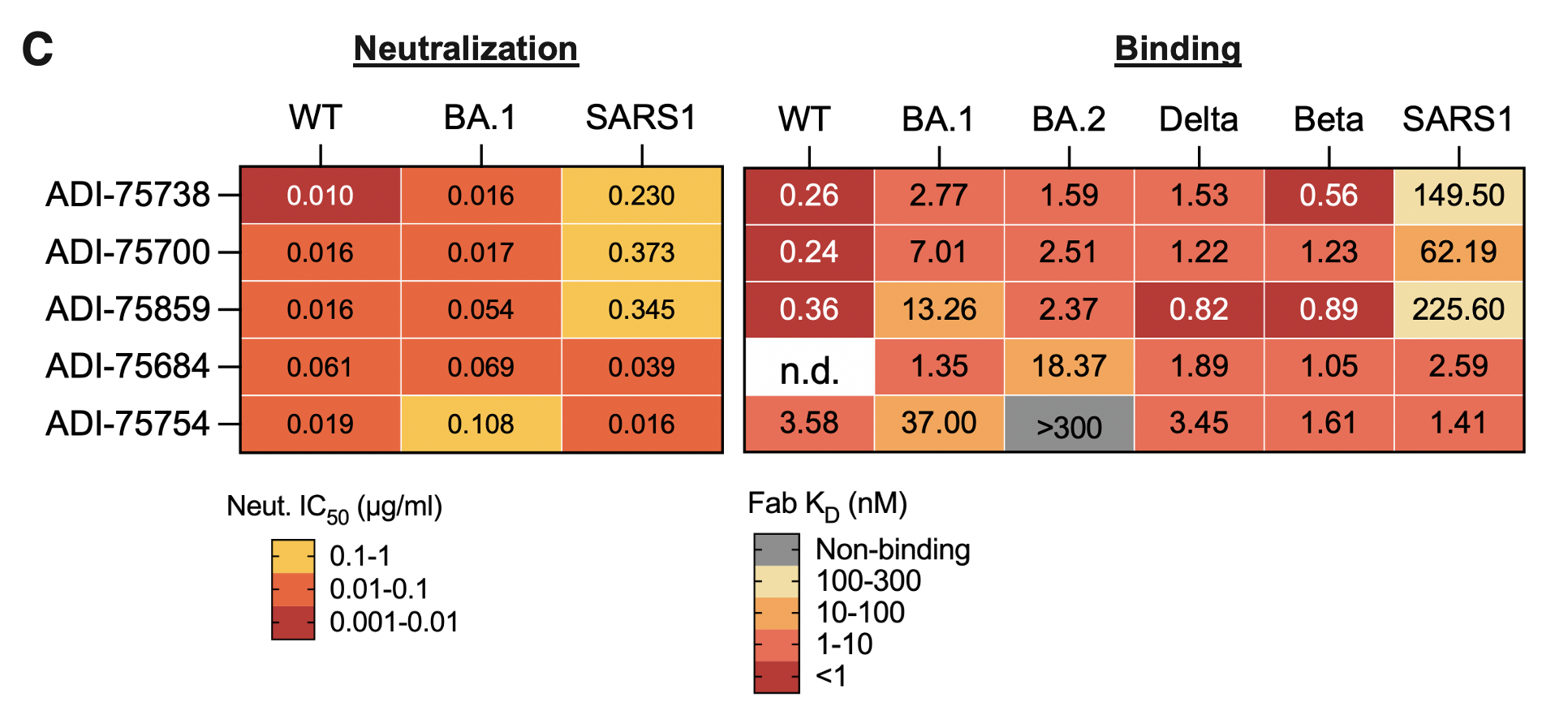

**Fig. S6.** Antibody breadth of activity against SARS-CoV-2 variants and SARS-CoV. **(A)** Fab binding affinities of BA.1 neutralizing antibodies to SARS-CoV RBD, as determined by BLI. (B) Neutralization IC_50_s against SARS-CoV and BA.1 among antibodies displaying Fab binding to SARS-CoV. (C) Neutralization potency (left) and Fab binding affinity (right) for the indicated SARS-CoV-2 variants of concern and SARS-CoV. IC_50_, 50% inhibitory concentration; KD, equilibrium dissociation constant; N.B., non-binding; n.d., not determined; N.N., non-neutralizing.

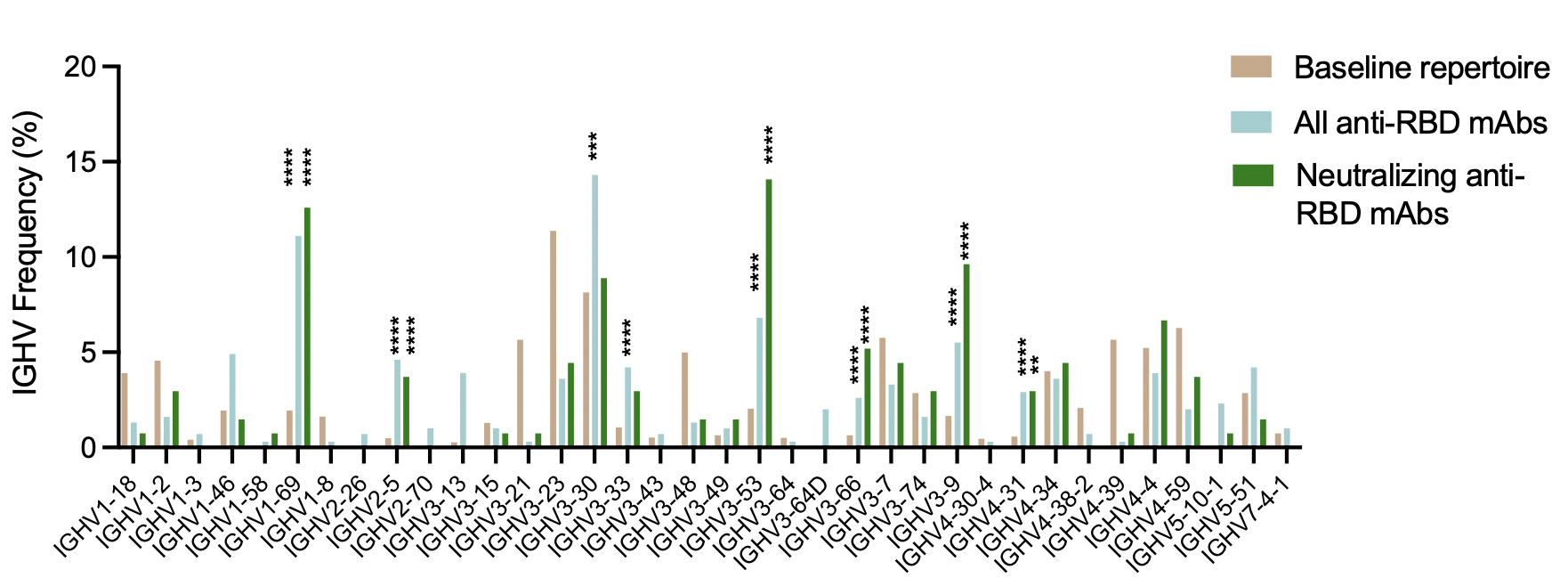

**Fig. S7.** IGHV germline gene frequency distributions among BA.1 neutralizing antibodies. Frequency distributions of all RBD-binding antibodies isolated from breakthrough infection donors and the human baseline (unselected) repertoire are included for reference (*17*). Statistical comparisons were made by Fisher's exact test compared to the baseline repertoire. IGHV, immunoglobulin heavy variable domain. *P < 0.05, **P < 0.01, ***P < 0.001, ****P < 0.0001.

.

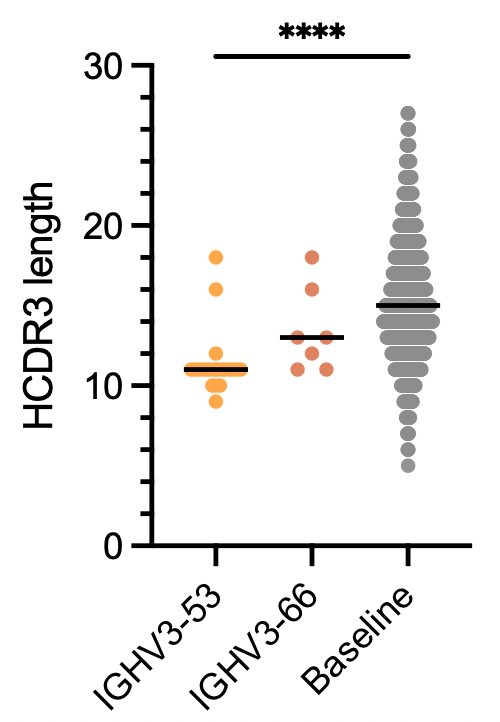

**Fig. S8.** HCDR3 amino acid lengths of BA.1 neutralizing antibodies utilizing IGH3-53/3-66 germline genes, with baseline antibody repertoire lengths included for comparison (*17*). Statistical comparisons were determined by Kruskal-Wallis test with subsequent Dunn's multiple comparisons. HCDR3, heavy complementarity determining region 3. ****P<0.0001

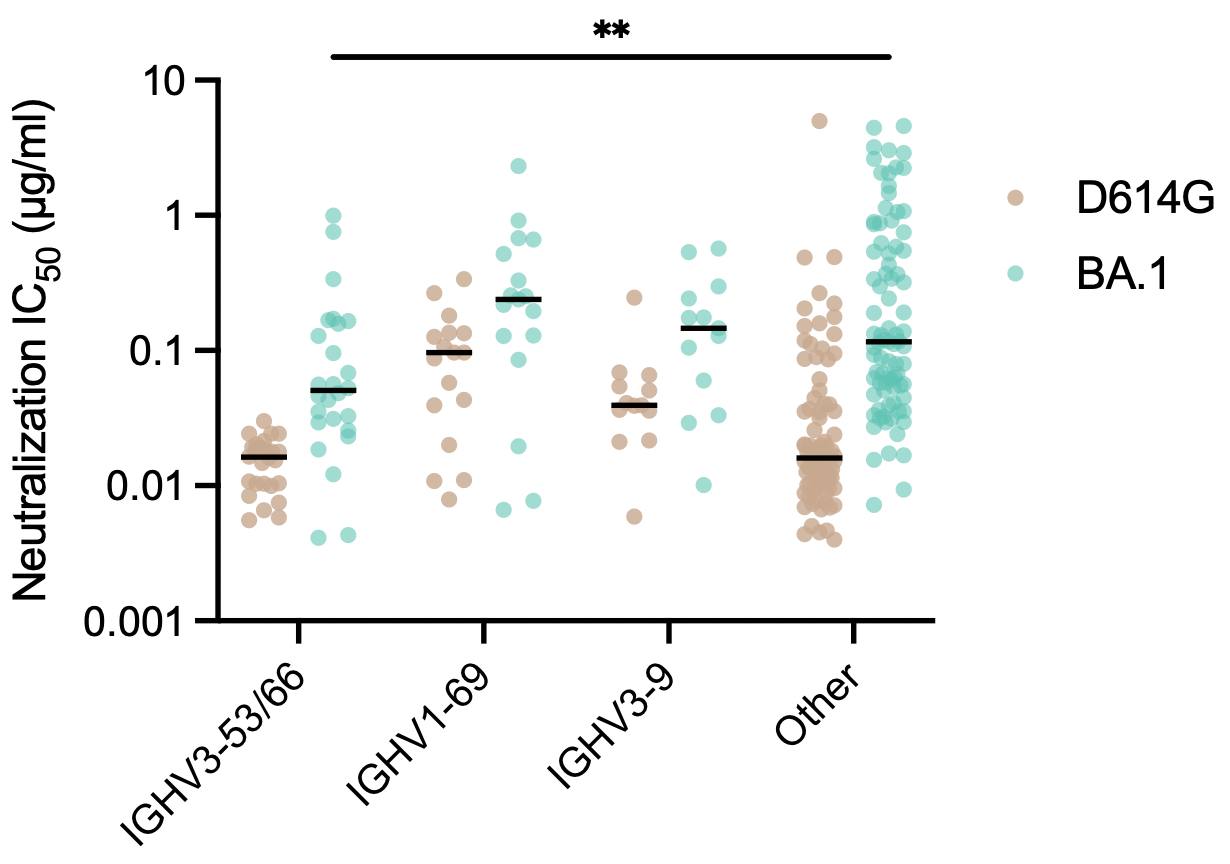

**Fig. S9.** Neutralizing activity of antibodies utilizing convergent IGHV germline genes against D614G and BA.1, as determined using an MLV-based pseudovirus assay. Black bars represent median IC_50_s. Statistical significance was determined by two-way ANOVA with subsequent Dunnett's multiple comparisons test IC_50_, 50% inhibitory concentration. IGHV, immunoglobulin heavy variable domain. **P < 0.01.

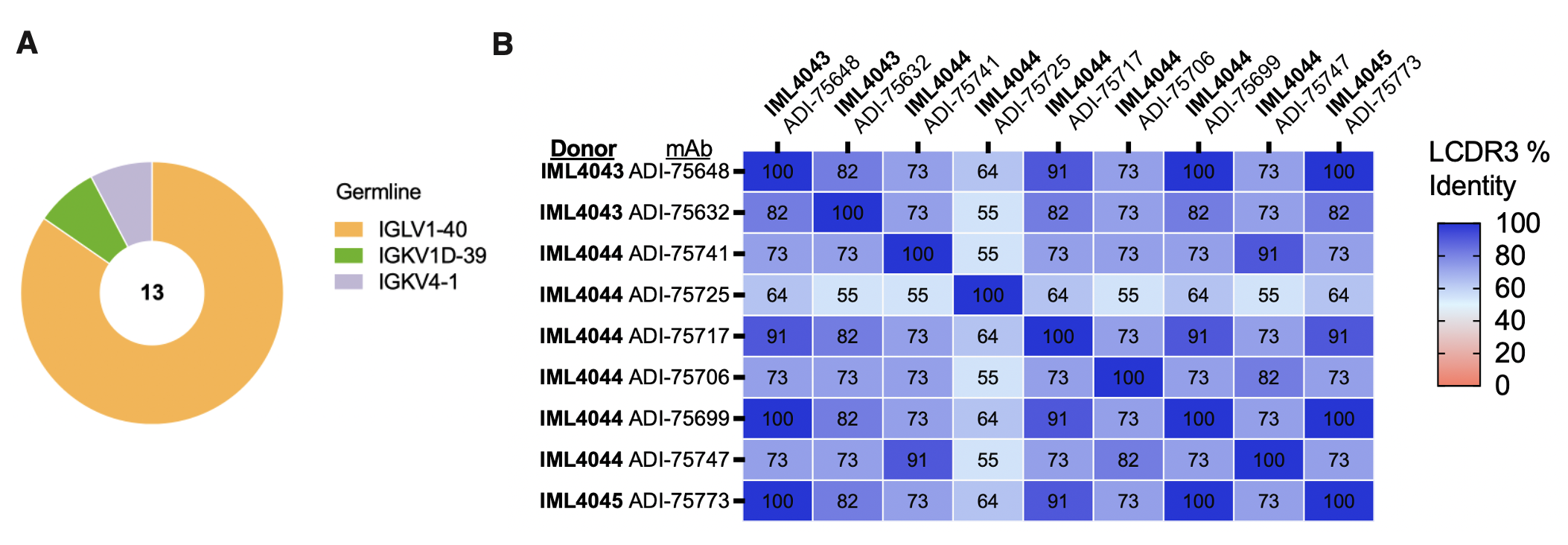

**Fig. S10.** Sequence features of breakthrough infection-derived IGHV1-69 neutralizing antibodies. (A) Light chain IGLV germline gene usage among non-ACE2-competitive IGHV1-19 antibodies. The total number of antibodies is shown in the center of the pie. (B) Pairwise comparisons of LCDR3 amino acid identity. The percentage identity of each pairwise comparison is shown in the middle of each square. LCDR3, light chain complementarity determining region 3; IGKV, immunoglobulin kappa variable domain. IGLV, immunoglobulin lambda variable domain.

**Table S1.** BA.1 breakthrough infection donor characteristics.

| **Donor ID** | **IML4041** | **IML4042** | **IML4043** | **IML4044** | **IML4045** | **IML4054** | **IML4055** |
| --- | --- | --- | --- | --- | --- | --- | --- |
| **Age** | 45 | 19 | 23 | 23 | 24 | 38 | 23 |
| **Sex** | F | F | M | F | F | F | F |
| **Vaccination History** | 2x BNT162b2 | 2x BNT162b2 | 2x BNT162b2 | 2x BNT162b2 | 2x BNT162b2, 1x mRNA-1273 | 2x mRNA-1273, 1x BNT162b2 | 3x BNT162b2 |
| **2nd dose vaccination date** | 7-May-21 | 22-Jul-21 | 23-May-21 | 10-Feb-21 | 15-May-21 | 5-May-21 | 1-May-21 |
| **3rd dose vaccination date (if applicable)** | - | - | - | - | 20-Dec-21 | 11-Dec-21 | 9-Dec-21 |
| **Date of PCR-confirmed infection** | 31-Dec-21 | 4-Jan-22 | 30-Dec-21 | 2-Jan-22 | 6-Jan-22 | 19-Jan-22 | 6-Jan-22 |
| **Days between breakthrough infection and sample collection** | 25 | 21 | 26 | 23 | 19 | 14 | 27 |

**Table S2.** Uninfected/mRNA-vaccinated cohort characteristics.

|  | **2x mRNA (1M)** | **2x mRNA (6M)** | **3x mRNA (1M)** |
| --- | --- | --- | --- |
| **Sample size** | 12 | 11 | 11 |
| **Age median (range)** | 31 (21-42) | 36 (26-42) | 54 (32-58) |
| **Vaccination regimens (%)** |  |  |  |
|  | 2x mRNA-1273 (75%) | 2x mRNA-1273 (82%) | 3x mRNA-1273 (64%) |
|  | 2x BNT162b2 (25%) | 2x BNT162b2 (18%) | 2x mRNA-1273, 1x BNT162b2 (36%) |
